# A Multiparametric Activity Profiling Platform for Neuron Disease Phenotyping and Drug Screening

**DOI:** 10.1101/2021.06.22.449195

**Authors:** Bruno Boivin, Kasper C.D. Roet, Xuan Huang, Kyle W. Karhohs, Mohammad H. Rohban, Jack Sandoe, Ole Wiskow, Rie Maeda, Alyssa Grantham, Mary K. Dornon, Jenny Shao, Devlin Frost, Dylan Baker, Kevin Eggan, Anne E. Carpenter, Clifford J. Woolf

**Author notes:** Corresponding author: Clifford J. Woolf. Co-first authors (Bruno Boivin, Kasper C.D. Roet).

## Abstract

Patient stem cell-derived models enable imaging of complex disease phenotypes and the development of scalable drug discovery platforms. Current preclinical methods for assessing cellular activity do not, however, capture the full intricacies of disease-induced disturbances, and instead typically focus on a single parameter, which impairs both the understanding of disease and the discovery of effective therapeutics. Here, we describe a cloud-based image processing and analysis platform that captures the intricate activity profile revealed by GCaMP fluorescent recordings of intracellular calcium changes and enables discovery of molecules that correct 153 parameters that define the amyotrophic lateral sclerosis motor neuron disease phenotype. In a high-throughput screen we identified compounds that revert the multiparametric disease profile to that found in healthy cells, a novel and robust measure of therapeutic potential quite distinct from unidimensional screening. This platform can guide the development of therapeutics that counteract the multifaceted pathological features of diseased cellular activity.

## INTRODUCTION

Neurodegenerative diseases are among the most difficult to treat. One such disease, amyotrophic lateral sclerosis, is associated with a progressive loss of upper and lower motor neurons leading to a gradual loss of control over the muscles vital for walking, talking, swallowing, and breathing, a debilitating and typically rapidly fatal outcome for patients. ALS remains a clinical challenge with only two FDA-approved drugs, both though with minimal life-prolonging effects. The prognosis for patients diagnosed with ALS is poor, with most patients succumbing to the disease within 3 to 5 years. The lack of effective treatments stems both from an incomplete understanding of the biological basis of the disease as well as the use of preclinical drug screening approaches that are not predictive of efficacy in patients.

In its familial form, ALS is caused by mutations in the *SOD1, C9orf72, FUS*, and *TARBP1* genes, among many others (*1*), and motor neuron electrical activity is severely impacted, as detected both in patients and patient-derived motor neurons (*2,3*). Mouse models as well as patient iPSC-derived motor neurons reveal alterations in the excitability phenotype across multiple parameters, including membrane potential, sodium and potassium peak currents, action potential firing, and synaptic activity (*4,5,6*). The development of therapies targeting aberrant electrophysiological properties usually focus only on a single electrophysiological property, the action potential firing rate (*7*). A recent study from our group successfully applied the approach of using a reduction in ALS human motor neuron firing rate to validate two known ALS activity modulators (Kv7.2/3 ion channels and AMPA receptors) and identify D2 dopamine receptors as a novel target (*8*). However, the excitability phenotype in ALS motor neurons is closely linked to multiple changes in cell health, such as ER-stress and unfolded protein response (*6*), and to changes in ion channel composition, molecular architecture and post-translational state, membrane trafficking, intra- and extracellular ion concentrations, and synapse strength, which impact diverse aspects of neuronal activity, especially in neurodegenerative disorders (*6*). We hypothesized that a multifaceted disease etiology requires a multiparameter analysis method and that simultaneous measurements of distinct cellular activity parameters such as peak amplitude, frequency, rise and fall time, and other kinetics, might provide a more comprehensive, and therefore superior basis, for defining complex disease phenotypes. In turn, multiparameter compound screening may also better detect disease-modifying actions of drugs than traditional single parameter-based assays. Molecules that counteract the disease pathology across multiple dimensions may provide superior rescue from a disease state and have a greater impact on disease progression than simply focusing on the restoration of a single feature, opening the possibility for novel and more precise therapeutics targeting the complexity of the disease state.

Here, we describe the development and application of a high-throughput cloud-based image processing and unbiased multiparametric activity profiling analytic platform. We call it CELLXPEDITE for its ability to process large volumes of cellular imaging data, thereby accelerating drug screening, and make it open source for use by the neuroscience community. By analyzing 153 parameters simultaneously we can automatically capture the complex activity profiles produced by diseased cells (ALS-patient derived carrying the *SOD1(A4V)* mutation) and the effects of candidate compounds on such cells, and from this identify compounds that convert the complex disease profile to one similar to that present in healthy cells. We show that such a platform can identify subtle perturbances in the activity of motor neurons and enables a robust selection of compounds that reverse the complex disease phenotype.

Most drugs for neurodegenerative diseases fail in clinical trials due to poor efficacy or unforeseen side effects, which is costly for the pharmaceutical industry and a health risk for the patients. Here we combine advances in human stem cell derived neuronal models, fluorescent reporters, high throughput live cell imaging systems, and cloud analysis platforms, to enable the discovery of molecules that can transform a complex disease phenotype to a healthy phenotype.

## RESULTS

### Extraction of Cellular Activity

During action potential firing, voltage-dependent calcium channels are activated and deactivated, leading to changes in intracellular calcium that allow measurement of neuronal activity through the genetically encoded calcium reporter GCaMP6 (*9*). We developed a robust high-throughput 384-well single-cell GCaMP6-based activity assay to record the spontaneous firing of iPSC-derived human motor neurons with and without the highly penetrant *SOD1(A4V)* mutation (Fig. 1A). To capture a multiparametric representation of GCaMP activity in each individual cell, we developed a scalable cloud-based live-cell image processing pipeline that automatically compensates for imaging artifacts, performs photobleaching corrections, identifies individual neurons, and extracts and denoises calcium transients (Fig. 1B,C). In contrast to existing commercial or open-source software packages (*10,11,12,13*), our workflow is seamless, extracts multiple complex parameters, and is suitable for large-scale drug screening. Our aim was to better capture the intricacies of cellular activity for disease modeling and enable high-throughput disease phenotype correction screening.

**Fig. 1.**
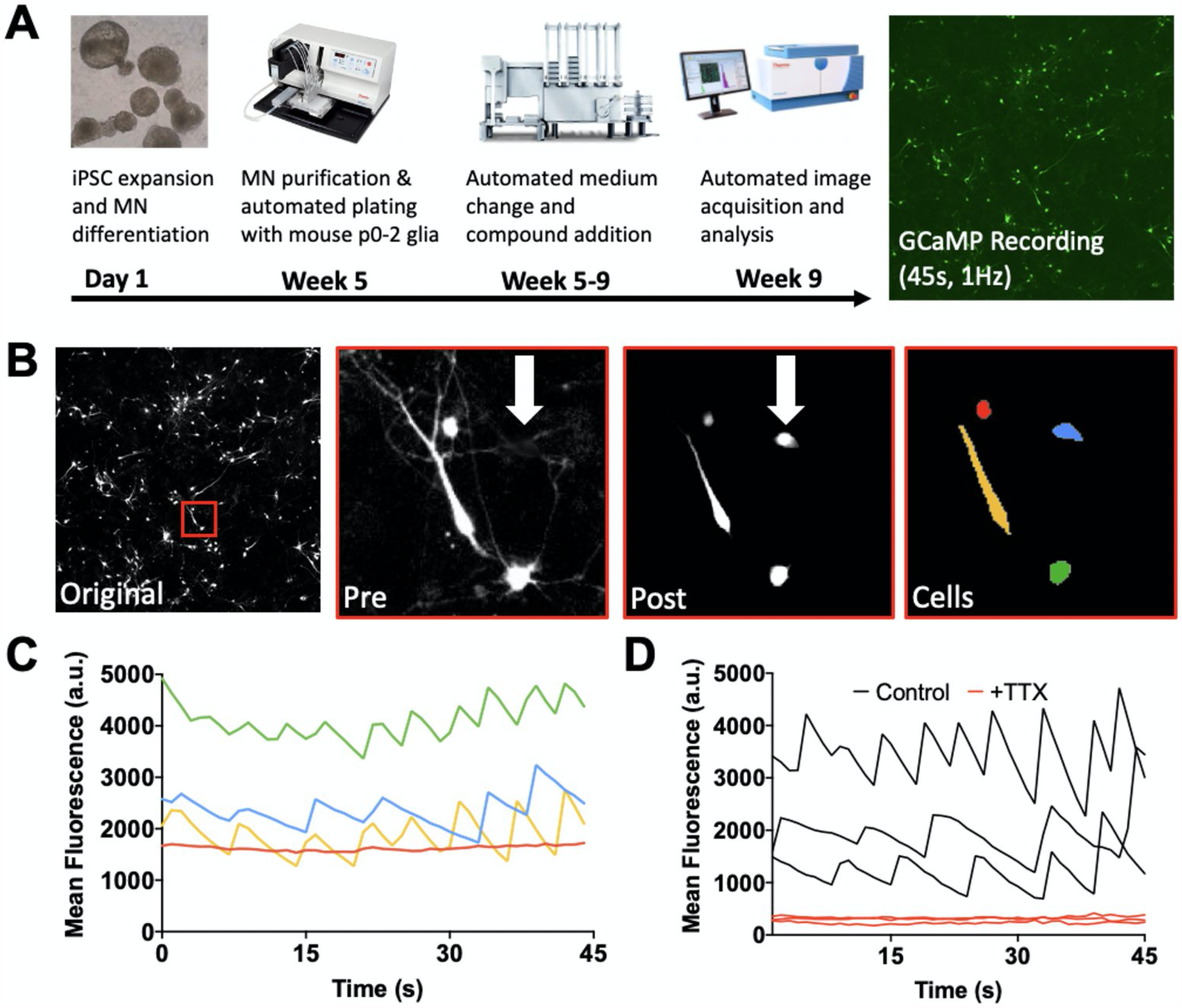
Cellular Activity Extraction Pipeline. **(A)** Schematic overview of the differentiation, plating, maturation, and imaging (whole field, 5x) of GCaMP6-positive human ALS patient-derived motor neurons. **(B)** Analysis of the fluorescence imaging data at a single time point does not allow for identification of inactive cells (“original” & “pre”). Temporal projection of the calcium imaging data over 45 seconds, combined with spatial filtering, enables identification of all cells regardless of activity (“post” & “cells”). **(C)** Fluorescence traces of cells identified in (B). **(D)** Spontaneous neuronal activity in dimethyl sulfoxide treated cells is eliminated by tetrodotoxin treatment (TTX).

High-throughput plate readers generally have an uneven spatial distribution of light across the field of view, which impacts both image segmentation and fluorescence quantification. We therefore incorporated into our pipeline modules from the open-source software CellProfiler (*14*) to perform illumination correction and enhance the visibility of neurons expressing GCaMP6 (Fig. S1). Our approach spatially resolved all cells, regardless of temporal activity, allowing us to determine the proportion of active cells and assess well contamination and compound toxicity. Calcium traces were automatically extracted for each neuron and compensated for background luminescence and photobleaching effects (Fig. S2). To validate that the residual signals were reflective of spiking activity, we extracted fluorescence traces in the presence and absence of tetrodotoxin (TTX), a sodium channel blocker that inhibits the firing of action potentials in neurons (*15*). TTX eliminated the spontaneous calcium wave fluctuations observed in control cells (DMSO vehicle) (Fig. 1D). Baseline fluorescence was also lower in the presence of TTX, suggesting that the measurements are sensitive to both absolute and relative differences in calcium levels.

### Multiparametric GCaMP Phenotypes

The comparison of GCaMP calcium transients between neurons, as well as changes due to drug action, has traditionally relied on counting the total number of peaks in fluorescence as the sole parameter (Fig. 2A), an extension of spike counting by electrophysiological recordings. However, because GCaMP6 fluorescence fluctuations are heterogenous and unevenly dispersed in time (Fig. 1C,D), simple metrics such as peak count or amplitude convey an incomplete picture (Fig. 2A). The disparity between the complex nature of cellular activity and its historical unidimensional characterization raises concerns over the accuracy of such measurements for reflecting the disease state of the neuron and its response to drug treatment.

**Fig. 2.**
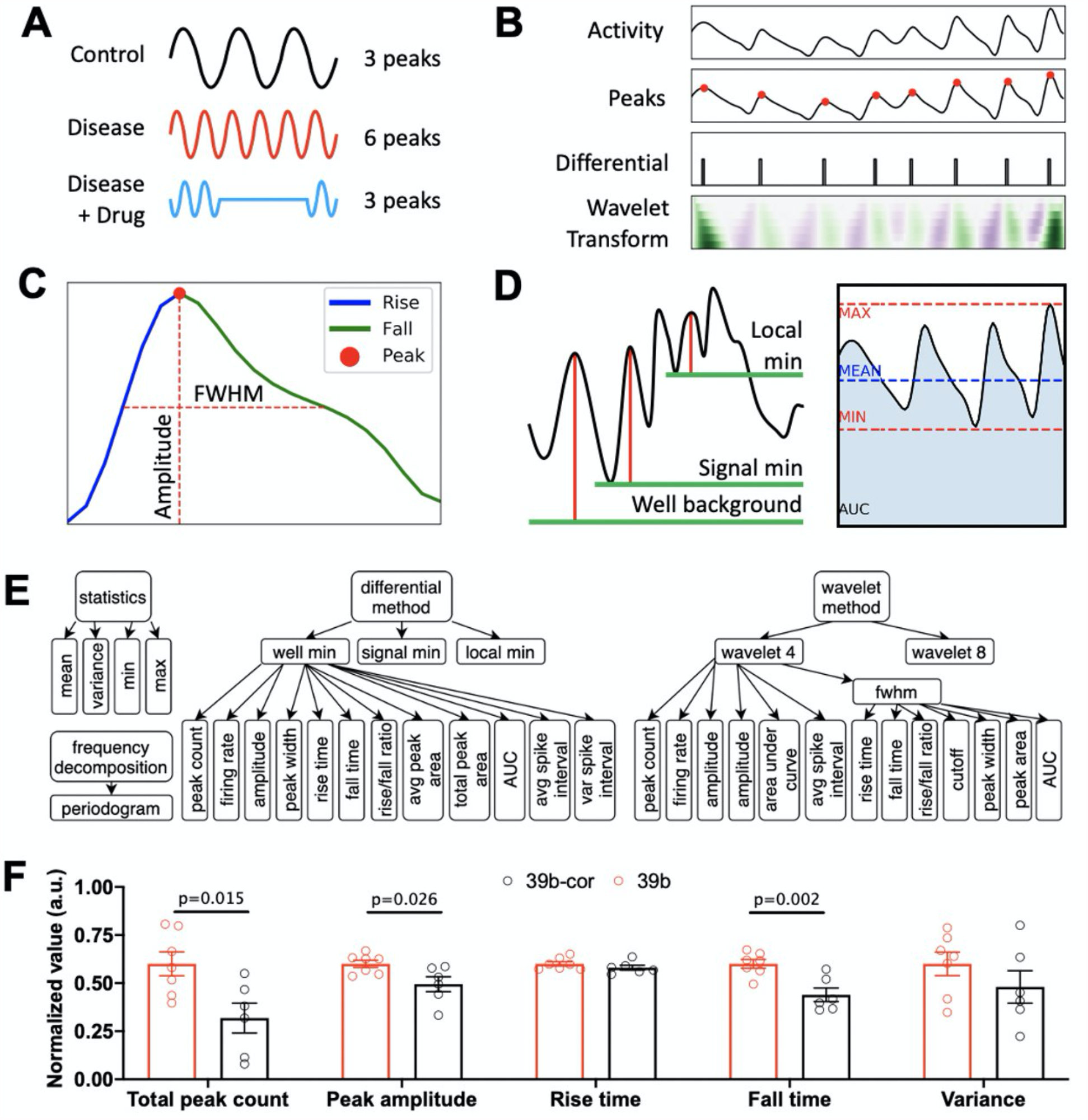
Parametrization of Cellular Activity. **(A)** Traditional peak counting does not identify temporal (e.g. uneven inter-spike intervals) or amplitude differences in calcium transients. **(B)** A combination of differential and continuous wavelet transforms accurately identifies peaks in activity. **(C)** Automated peak deconvolution quantifies the rise, apex, fall, amplitude, and full width at half maximum of an individual peak. **(D)** Cellular peak parameters are quantified relative to three baselines: well background intensity, cell minimum intensity, and intensity at the peak onset (left). Signal-wide features of activity, including dispersion metrics (minimum, maximum, mean) and the area under the curve (AUC), are automatically captured (right). **(E)** Breakdown of the 153 activity parameters computed for each cell. A single node under the differential and wavelet methods is expanded for clarity; the other nodes share the same parameter subtree. **(F)** Spontaneous GCaMP activity analysis of SOD1^A4V^ human motor neurons compared to isogenic corrected motor neurons (39b-cor) reveal variable differences across five example parameters. Features are normalized to the average of the 39b group and presented as mean ± SEM using Welch’s t-test.

To provide a more comprehensive readout of excitability in diseased and healthy motor neurons, we set out to parametrize spontaneous fluctuations of the calcium reporter signal. We identified peaks in activity using a differential approach with high sensitivity to rapid fluctuations as well as continuous wavelet transforms (*16*) to account for multi-scale peaks (Fig. 2B). Peaks were then individually parsed to quantify subtle kinetic differences and fluorescence shifts (Fig. 2C). Relative and absolute changes were detected by establishing 3 baselines for the measurements, namely the well background intensity, the cell minimum intensity, and the intensity at peak onset (Fig. 2D). To complement the individual peak dissection with signal-wide features, statistical dispersion metrics were computed along with the area under each signal (Fig. 2D). Finally, the power spectrum of the calcium traces, i.e. the distribution of power into frequency components composing the signals, were obtained by computing their discrete Fourier transform. A total of 153 parameters spanning both time and frequency domains were acquired to describe the activity of each cell (Fig. 2E), providing a comprehensive breakdown of activity, and widening the scope of the activity phenotype across multiple axes.

### Healthy and Disease Activity Profiles

To characterize the complex ALS disease phenotype, we utilized the image processing and analysis platform to generate and study all the parameters derived from the spontaneous activity of two cell lines derived from a human ALS patient (39b: ALS disease model and 39b-cor: isogenic corrected healthy control). In addition to the expected increased peak counts in the disease cell line, we discovered changes in activity parameters reflective of slower kinetics and higher peak amplitudes (Fig. 2F). With the demonstration that cellular activity is best captured using multiple parameters, we then conducted a full multiparametric analysis of the disease (39b) and control (39b-cor) motor neuron activity phenotypes. These phenotypes were generated by averaging the multiparametric activity profiles across 8 replicates for each cell line. From these profiles, we calculated the absolute difference across all parameters between the two cell lines, which resulted in a disease profile consisting of 153 activity parameters (top 50 distinguishing features presented in Fig. 3B). Surprisingly, the largest differences were found in features related to peak amplitude as opposed to peak count, though the latter remained in the top 5 distinguishing features. We found properties from the frequency domain to be of lesser importance. The multiparametric disease profile was then used to identify phenotypic rescue of all the disease distinguishing cellular activity parameters back to those in the healthy control.

**Fig. 3.**
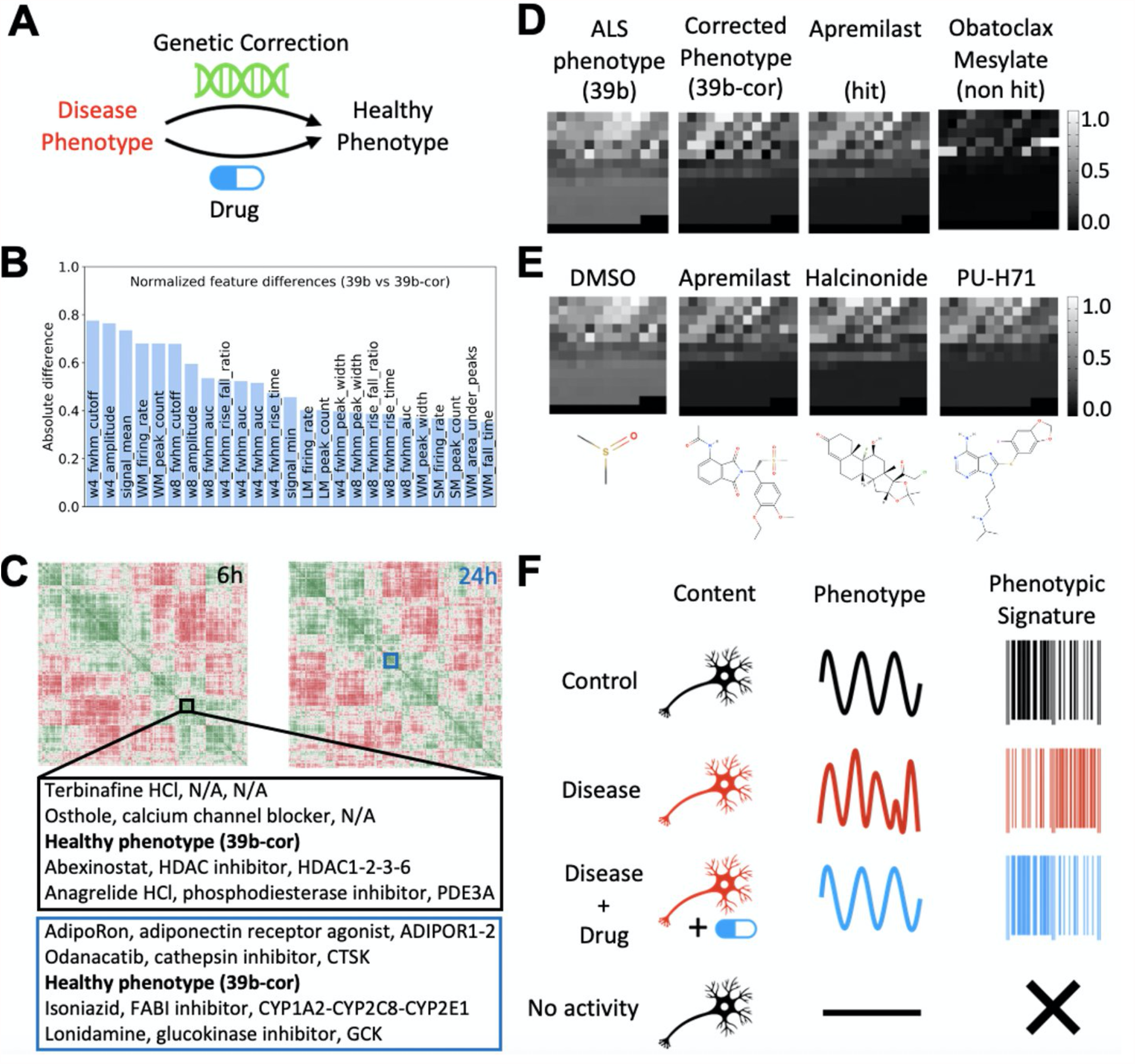
Multiparametric Drug Screening Strategy. **(A)** Schematic illustrating that both genetic mutation correction and drug action can restore a healthy activity phenotype in a diseased cell line. **(B)** Ranking of the top 25 individual parameter differences between the disease (39b) and corrected cell line (39b-cor) reveal relative importance of features. **(C)** Hierarchically-clustered heatmap of 1902 compounds from the Selleck bioactive library 6h and 24h after treatment; each compound is represented using the 153 activity features normalized to the control phenotype. Compound name, mechanism of action, and molecular targets of the 4 hits that resulted in cellular activity closest to the healthy phenotype at each timepoint is shown. **(D)** Comparison of the multiparametric activity signatures for the disease phenotype (39b), the healthy phenotype (39b-corrected), a hit compound (Apremilast) and a non-hit compound (Obatoclax Mesylate) on 39b MN distinguishes activity patterns. Features are ordered based on preassigned positions in the grid and normalized to fit on the same intensity scale. **(E)** Comparison of the negative control (DMSO) and 3 hits (Apremilast, Halcinonide, PU-H71) that replicated in a validation screen on 39b MN reveals similar activity profiles and distinct molecular structures across the hits. **(F)** Depiction of phenotypic signatures capturing complex activity phenotypes in control and disease neuronal populations (first two rows). The action of an effective drug on diseased cells restores the control (healthy) phenotype (third row). No phenotypic signatures are produced for inactive cells (fourth row). Comparison of signatures provide a basis for measuring drug efficacy.

### ALS Multiparametric Drug Screen

Given the rich multiparametric readout of the ALS hyperexcitability phenotype, we sought to identify disease correcting drug candidates based on their specific effects on disease activity profiles. We screened 1902 compounds from the Selleck bioactive compound library, which includes FDA-approved compounds, active pharmaceutical agents, chemotherapeutic agents, and natural products. Compounds were mapped to points in the multidimensional space defined by the 153 features in their activity profiles (Fig. 2E), with distances between them reflecting the degree of similarity between the phenotypes they produced. We discovered a cluster of compounds which caused the disease-allele-containing cells to change and resemble control healthy cells, providing a pool of promising disease-correcting candidates (Fig. 3C). Among these were phosphodiesterase inhibitors and inhibitors of the mTOR pathway, both previously associated with neuroprotection (*17,18*). Because our pilot screen suggested that different activity profiles between the disease and control states reflect the modulation of activity, as opposed to a complete blockade of activity, we also investigated the effects produced by calcium-channel blockers which abolish the GCaMP signals triggered by action potentials. While these blockers clustered together, they were separated from the control phenotype and were not disease correcting, highlighting the sensitivity of the multiparametric approach to discriminate distinct activity states.

After excluding compounds that resulted in potent inhibition of neuronal activity (including for example, 5 µM tetrodotoxin, see table S1), we selected the 65 compounds with the shortest distance (strongest similarity) to the control profile, and tested an additional 6 replicates for each candidate over two independent recording sessions. We advanced the 80% most-consistent compounds (Fig. S3C) for further analysis. After ordering these compounds by their ability to normalize the disease state back to a healthy phenotype, as measured by the distance between the multiparametric profiles, the three compounds that most strongly and reproducibly normalized activity in the disease motor neurons were apremilast CC-10004 (phosphodiesterase inhibitor), halcinonide (glucocorticoid receptor agonist), and PU-H71 (a heat shock protein inhibitor).

Next, we wanted to confirm the benefits of using a multiparametric method over the traditional unidimensional approach. In addition to the 48 compounds that abolished neuronal activity (Table S1), i.e. compounds that resulted in complete activity suppression as opposed to a normalization of the disease activity profile, we also identified hits which only reduced the total number of calcium transient peaks, the top three of which included dexmedetomidine (adrenergic receptor agonist), bimatoprost (prostanoid receptor agonist), and tianeptine (atypical antidepressant). The hit compounds obtained using the multiparametric approach were approximately 20% closer to the targeted healthy control than those obtained from a one-dimensional peak count metric. These results demonstrate that utilization of multiparametric selection criteria offers superior specificity by imposing strict constraints, reducing the hits to only those that reverse the disease phenotype across all 153 dimensions. Therefore, screening against multiple disease-related parameters, as opposed to a single one, is likely to lead to the identification of completely different sets of compounds with distinct mechanisms of action and molecular targets, and which are therefore likely to have quite different efficacy profiles for the disease in patients.

### Activity Fingerprints

Next, we compared the hit compounds by converting their activity profiles into quantifiable visual representations (phenotypic signatures), negating the need to perform dimensionality reduction by providing a lens into the multidimensional activity space. We found the activity signatures of the ALS disease phenotype and the healthy phenotype to be quite different, while the signatures from compounds that closely matched the control phenotype were similar to each other and to the healthy profile (Fig. 3D). Compounds that introduced changes distinct from the disease or healthy state (potential undesired effects) had very different signatures (Fig. 3D). Interestingly, while the activity signatures of the disease-phenotype-correcting hits were comparable, their molecular structures and targets were different (Fig. 3E). These results suggest either that multiple different molecular mechanisms can be utilized to normalize pathological excitability profiles in SOD1^A4V^ motor neurons or that the off-target effects of these quite distinct compounds might be similar, although the latter is statistically most unlikely.

## DISCUSSION

We have designed a computational approach that combines the temporal and spatial resolution of high content microscopy imaging and automated high-throughput screening with the scalability of cloud computing, to accelerate the search for new molecules capable of rescuing an ALS motor neuron disease phenotype. The pipeline processes imaging data from individual wells in parallel to maximize throughput. Imaging artifacts are removed by a multi-layered correction algorithm and objects are identified by CellProfiler modules combined with a temporal profile analysis. While we collect information about the morphology, eccentricity, and surface area of cells, our analytic approach focuses on the dissection of temporal dynamics of GCaMP activity. The calcium dynamics encoded in each time series is parameterized into 153 features that are normalized and combined to form an overall representation of cellular activity (an activity fingerprint or profile). The generation of a multiparametric disease profile combined with unbiased hierarchical clustering of drug efficacy over 153 parameters simultaneously, allowed us to identify drugs that can convert a disease profile into a healthy profile, and also identify potential undesired side effects, from the perspective of a distortion in the multidimensional representation of healthy activity.

Through our large-scale bioactive compound drug disease phenotype correcting screen, we identified Apremilast, Halcinonide and PU-H71 as compounds that normalize the multifaceted GCaMP-based disease activity phenotype back to a healthy phenotype, providing a proof-of-principle for a new drug screening paradigm for ALS, and other diseases, though future study is needed to identify the molecular targets underlying these three compound’s actions. Although there are similarities in the chemical structures of Apremilast, Halcinonide and PU-H71, they are not members of the same class of molecules and have different annotated molecular targets, specifically phosphodiesterase 4, corticosteroid hormone receptor and heat shock protein 90, respectively. It is therefore possible or even likely that the observed drug effects are the result of interactions with different targets, thus any structure-activity relationship (SAR) campaign following this kind of multidimensional phenotypic screen will have to embrace the complexity of the disease and include a multi-target deconvolution process. This strategy may be a necessity for the development of successful therapeutics for complex neurological disorders such as ALS and represents a move from single target identification.

The ability to generate complex stem cell derived neuronal models for diseases such as ALS has provided a powerful tool to understand the known genetic forms of the disease. However, for most ALS patients the cause of their disease is unknown, and they are classified as sporadic. The development and execution of an automated and unbiased high throughput multiparameter activity profiling platform provides an unprecedented opportunity to interrogate neurons derived from many sporadic patient stem cell lines. Such an interrogation could lead to the identification of patients with a shared abnormal activity profile, and in this way, aid our understanding of common complex changes in neuronal activity in disease conditions, as well as the discovery of compounds that can rescue a comprehensive disease phenotype to a healthy one.

We evaluated the performance of our computational approach in terms of its usability, speed, and accuracy. In comparison to previous approaches, the automated measurement of 153 features of activity for each cell removes the inherent bias associated with manual selection of a subset of features and provides a highly sensitive framework for identifying activity disturbances. From a usability standpoint, this negates the need for user-specified parameters that are often arbitrarily chosen and biased. Additionally, our cloud-based design can theoretically scale to any number of wells provided sufficient computing resources are available. Since wells are processed in parallel and the results combined in constant time, increasing the number of compounds in a drug screen only marginally increases total processing time. Prior assays that include manual interventions are time-consuming, error-prone, and often not reproducible. Once adjusted for the type of cells being studied and the recording frequency, our approach does not require human intervention and can be applied to any number of recordings, provides comprehensive activity profiles, and is entirely reproducible. This is particularly useful when studying complex genetic diseases such as ALS, since it allows for the comparison of the activity profiles of different mutations under the same treatment. Therefore, this computationally efficient and deterministic approach, one that can be launched by the push of a button, is we argue, superior to previous methods in many regards.

In principle, our approach can be used with any cell, for any cellular functional activity that can be fluorescently tagged, and for which dynamics is a biologically relevant property. Because it is readily deployable on cloud infrastructures, it provides a seamless way to interrogate disease activity and the corrections in this phenotype introduced by drug actions. However, the reliance on dynamic data alone excludes morphological changes such as cell size and dendritic arbor changes, which may be important for disease progression. Previous studies, as well as our own observations, indicate that the soma size of motor neurons carrying the *SOD1*^*A4V*^ mutation is altered as the disease progresses (*2, 19*). A next logical step for the platform would be to combine activity-based phenotypes with detection of morphological aberrations.

We anticipate that sophisticated phenotypic screens that embrace the complexity of cellular function and delineate multiple facets of drug action will accelerate the discovery of successful and novel therapeutic interventions. Our findings demonstrate the existence of multiple complex and previously unexploited facets of the ALS excitability phenotype and the discovery of compounds that robustly correct many parameters of this phenotype simultaneously. This discovery, coupled with the sensitive cloud-based high-throughput multiparametric framework for capturing and analyzing subtle changes in cellular activity, represents a substantial leap forward for comprehensive disease profiling and drug screening in the pursuit of novel treatments for ALS and other diseases. Improvements to the earliest phases of the translational process, which are inextricably linked to success of clinical trials, have the potential to both significantly reduce costs and health risks for patients as well as improve efficacy. Ultimately, the combination of human stem cell models and novel multiparametric screening approaches have, we argue, the promise to lead to safer and more effective therapeutics for ALS.

## MATERIALS AND METHODS

### Single Cell GCaMP6 Screen

iPSC’s were expanded and differentiated into motor neurons using an embryonic body-based (EB) protocol as previously described (*2*). At day 24, EBs were dissociated to single cells with accutase, frozen and stored in liquid nitrogen until FACS purification of motor neurons that were NCAM (BD Biosciences, #561903) positive and EpCAM (BD Biosciences, #347198) negative. Motor neurons were co-cultured in Greiner Bio-one cell culture plates 384-well plates at a density of 7,500 cell per well with 10,000 glial cells obtained from P0–P2 C57BL/6 pups (Jackson Laboratory) as described in Di Giorgio (*20*). Mouse procedures were approved by Boston Children’s Hospital Institutional Animal Care and Use Committee (IACUC).

Cells were plated in the presence of 10 µM ROCK inhibitor Y-27632 (Tocris) and 1µM EDU (Thermo) in motor neuron media (Neurobasal media) (NB, Invitrogen, Carlsbad, CA), supplemented with B27, N2 supplement, glutamax and non-essential amino acids (GIBCO™, Thermo Fisher), 10 ng/ml each of BDNF, GDNF, and CNTF (R&D Systems, Minneapolis, MN), and 0.2 mg/ml ascorbic acid (Sigma). After incubation for 2 days, media was replaced every 3 or 4 days by Neurobasal media. LV-synapsin-GCaMP6 viral infections were performed in weeks 2 and 3 at a MOI of 5. After 3-4 weeks, synaptic blockers are added to each well at the following concentrations: Bicuculline 25µM, Strychnine 10µM, AP-5 100µM, CNQX 10µM. After 2hr chemogenomic library compounds from the Selleck Bioactive Compound Library (plates 3651-3656; ICCB Longwood Screening Facility) were added at a final concentration of 3µM. DMSO was used as a negative control at 0.03%. Tetrodotoxin 5µM in 0.03% DMSO was used as a positive control. After 6hr and 24hr, the plates were recorded for 4-5hr in the Arrayscan™ XTI (Thermo Fisher) with excitation at 485 nm and emission at 521 nm. Recordings were acquired at 1Hz for 45 seconds over two independent sessions.

### Illumination Correction

Uneven illumination was corrected across the field of view using a CellProfiler (*14*) pipeline that sequentially applies two built-in modules. The module *CorrectIllumCalculate* was first included to calculate the illumination function from all images across cycles. The smoothing method was set to *Median Filter* with default settings to capture global illumination disparities rather than cell-specific variations in brightness. The other parameters of the module were left unchanged. The *SaveImages* module was then added to save the output upon the last cycle as a 32-bit floating-point tiff image.

### Maximum Projection

The maximum value of each pixel across all frames in the recording was computed and saved as an 8-bit frame representing the maximum projection of the recording using the Python programming language (*21*).

### Cellular Fragments Identification

Cells were first identified and morphologically characterized from the maximum projection image by applying a CellProfiler (*14*) pipeline that combines thirteen sequential modules. A sample maximum projection subimage is followed through each module in Figure S1 B. Starting with the original image (b1), a median filter is applied to blur away small artifacts, with the typical artifact parameter set to 3 (b2). Neurites are then computationally enhanced to become more visible and identifiable using the *EnhanceOrSuppressFeatures* module, with the tubeness enhancement method and smoothing scale set to 2 (b3). The *ImageMath*module is used to average the enhanced features with the blurred image, generating a cleaner delineation of the contours of interest (b4). Clustering-based image thresholding is performed using the Otsu’s method to separate foreground from background pixels using the *IdentifyPrimaryObjects* module (b5). Simultaneously, objects with diameters ranging from 5 to 35 pixels are identified and their contours outlined (b6). The *IdentifySecondaryObjects* module is used to identify secondary objects such as neurites using the propagation method, which helps find dividing lines between clumped objects and refine cell segregation (b7). The brightness, morphology, and texture of individual fragments is characterized using a combination of modules, specifically *MeasureImageIntensity, MeasureObjectIntensity, MeasureObjectSizeShape*, and *MeasureTexture*. Consolidation of the location of all pixels belonging to cellular fragments generate a numerical mask saved as a 16-bit tiff image which can be used to identify all or specific subsets of cells (b8). Background pixels are assigned a value of 0 while other pixels are assigned a unique fragment-specific integer identifier between 1 and K, where K is the total number of fragments identified in the well.

### Spatial Proximity Assessment

Pairwise spatial proximity of fragments is assessed by measuring the Euclidean distance between their centroids. Centroids are calculated using Python’s SciPy library. This generates an NxN matrix, where N is the number of fragments and each entry in the matrix the distance in pixels between two non-overlapping fragments.

### Activity Similarity Assessment

Pairwise time series similarity is assessed using the standard implementation of the dynamic time warping algorithm included in the *mlpy* (*22*) Python library with default parameters. An unnormalized minimum-distance warp path is computed for each pair of signals. An NxN matrix is thus constructed, where N is the number of fragments and each entry in the matrix the cost associated to the warping path between the signals of two non-overlapping fragments.

### Cellular Fragments Consolidation

The matrices described in Figure S1 D are first normalized to their respective means through division. The geometric mean of their normalized forms is then computed from the Python implementation included in the *scipy*.*stats* library (SciPy version 1.2.1) (*23*), yielding a matrix with each entry being a unified distance between each pair of fragments. The resulting matrix is fed into an agglomerative hierarchical clustering algorithm (*24*) that utilizes complete linkage to merge the subtrees. More specifically, the *scipy*.*cluster*.*hierarchy* library is used to compute the linkage matrix and to retrieve flat clusters from it, with a threshold of 0.1. The threshold corresponds to the height at which the dendrogram resulting from the hierarchical clustering is cut. Branches below the cut threshold are pruned, yielding consolidated fragments. Branches above the threshold are deemed part of distinct cells. The cell mask previously generated is updated so that pixels belonging to the same cell are assigned the same unique integer identifier. The identifiers assigned range from 1 to N, where N is the total number of cells after the fragments have been recombined.

### Extraction of Cellular Activity

From the whole-cell mask obtained by combining cellular fragments, a mapping from unique cell identifiers to their respective set of pixels is constructed. The activity of individual cells is extracted by averaging at each time point the brightness of their respective pixels.

### Decay Correction

The decay in fluorescence is estimated by averaging the decay of randomly selected pixels from each well. Specifically, 100 pixels from each well were randomly selected and grouped. A first-degree exponential decay function was fitted to the aggregated data using the *scipy* curve fitting module, with the model function *y* = *ae*^−*bx*^ + *c*, the bounds constricted to the positive range, and an initial guess of the parameters of *a* = 2000, *b* = 0.01, and *c* = 200. The resulting fit curve is normalized through division by its maximum value. The activity of each cell over time is then divided by this normalized decay curve.

### Background Subtraction

Using the cell mask to identify background pixels, a sample of 10,000 background pixels are randomly selected and their intensity averaged at each time point. This generates a time series that estimates background intensity over time. This curve is then subtracted from the activity of each cell.

### Peak Identification

Peaks in cellular activity were identified using code derived from the *find_peak* function of the *scipy* package. Peaks were labeled on their rising edge with a minimum interpeak distance of 1. Troughs were identified by applying the same approach onto the inverse of the time series. Peaks were also identified using the continuous wavelet transform method provided in the *scipy* package. Wavelets of width up to 4 (W4) and up to 8 (W8) were used separately to identify distinct sets of peaks, each time with a gap threshold of 3 and a maximum distance of half the maximum width. The W4 approach used a minimum ridge line length of 3, a minimum SNR ratio of 1, and a noise percentile of 10. The W8 approach used a minimum ridge line length of 5, a minimum SNR ratio of 2, and a noise percentile of 10.

### Baselines

Three baselines were established to compute individual peak characteristics. Since background intensity was subtracted from each signal, measurements relative to the y=0 line were used to obtain features relative to the background intensity. The respective minimum fluorescence value of each cell across the 45-second recording was used a second baseline. The third baseline was determined for each peak by taking the fluorescence value at the onset of the peak.

### Peak Threshold

For each baseline, a peak threshold was computed to distinguish noise from actual peaks. The noise level was established by taking the amplitude distribution of all peaks in the positive control wells, here those treated with tetrodotoxin. The spontaneous peak level was established by taking the amplitude distribution of all peaks in the negative control wells, here those treated with dimethyl sulfoxide (DMSO). The peak threshold was then calculated using the Otsu method from the combined distribution of peak heights using the SciPy Python package.

### Peak Features

Under the differential method, for each peak and each baseline, amplitude was measured relative to the baseline. Peaks whose height fell below the minimum peak threshold were discarded. Peak count was computed as the remaining number of peaks. Rise time was calculated as the time between the peak apex and the previous trough. Fall time was calculated as the time between the peak apex and the following trough. Peak width was measured as the time between the onset of the rise and the end of the fall. The area under a given peak was computed using the trapezoidal rule along the width of the peak. The average and variance of the time interval between peaks was also calculated.

Under the wavelet methods, the full width at half maximum was used as a reference point for computing the peak features. Rise time was calculated as the time between the half-maximum point along the rise and the peak apex. Fall time was calculated as the time between the peak apex and the half-maximum point along the fall. The other peak features were obtained using the same logic used under the differential method.

### Signal Features

Statistical metrics including the minimum, maximum, mean, and variance of each trace were computed using the *numpy* (*25*) package (version 1.18) in Python. The area under the entire signal was approximated using the trapezoidal rule provided in the same package. The power spectral density of each trace was obtained using the *periodogram* module of the *scipy*.*signal* package.

### Multidimensional Activity Clustering

Individual cell activity is described by the set of 153 features. Individual well activity is computed as the mean of each feature across all cells in the well. The control phenotype is obtained by averaging the features from the control wells, here the healthy control line (39b-cor). The disease activity phenotype is obtained by averaging the features from the disease wells, here ALS (39b). For each well, the features were normalized to the control phenotype features using Cytominer (*26*), which normally distributes the data around the control phenotype. Pairwise comparison of well activity phenotypes is evaluated by measuring the Euclidean distance between the wells in the multidimensional feature space using the SciPy spatial module. The resulting matrix is plotted as a hierarchically clustered heatmap using *seaborn* (*27*) (version 0.9.0). Compounds that result in similar activity phenotypes are observed as clusters in the heatmap. From the pairwise distance matrix, wells with activity phenotypes most similar to the control phenotype are extracted and ranked.

### Activity Signatures

For each well, each feature is normalized to the [0;1] range based on all observed values. The normalized feature vector representing each well is converted into a matrix of size 13×13 using the Python package *numpy*. The first 153 entries in the matrix correspond to the normalized features; the remaining entries are set to 0. The resulting matrix is used as the well activity signature and visualized in the form of a color-coded grid generated using the Python package *matplotlib* (*28*). Grid squares are colored based on a grayscale gradient where 0 is black and 1 is white. The position of each feature in the grid is preassigned to allow direct comparison of signatures.

### CELLXPEDITE Platform

We have released our implementation of the cloud-based processing and analysis platform as an open-source software (https://github.com/brunoboivin/cellxpedite). It is currently designed for deployment on Amazon Web Services, but it is also compatible with local clusters and other cloud computing service providers with minimal changes to the deployment scripts.

## Acknowledgments / Funding

This project was funded by grants from the Defense Advanced Research Projects Agency HR0011-19-2-0022, NIH R35NS105076 (CJW), Target ALS, a Milton Safenowitz Post Doctoral Fellowship for ALS Research, and a Boston Children’s Hospital-Broad Institute Collaboration Grant.

## Author contributions

B.B., K.C.D.R., X.H., C.J.W. wrote the manuscript. B.B., K.W.K., M.H.R. designed the analytical approach and wrote software for processing calcium imaging data. K.C.D.R., X.H., R.M., A.G., M.K.D., D.F., J.S., O.W., D.B, J.S. designed and performed the biological experiments. K.C.D.R., C.J.W., K.E and A.C. designed and supervised the study.

## Supplementary Materials

**Fig. S1.**
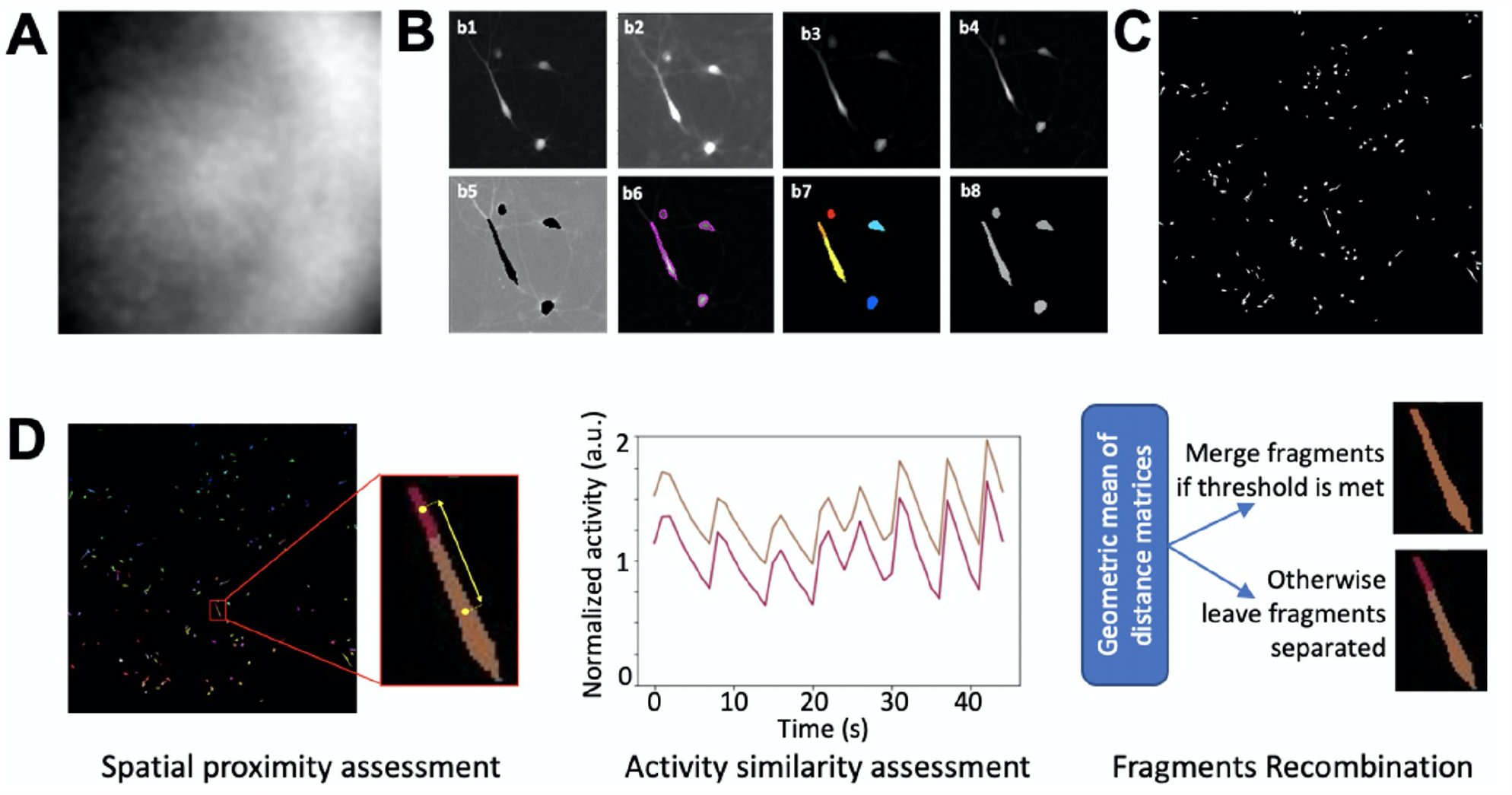
Cell Identification Strategy. **(A)** Uneven illumination captured using our pipeline. **(B)** Cell identification workflow applied onto a portion of the field of view. Starting from the maximum projection of the original recording (b1), a median blur is applied (b2) and combined with enhanced neurite features (b3) through averaging (b4). Background/foreground pixels are distinguished (b5), primary cell contours outlined (b6), secondary clumped cells separated (b7), ultimately producing a mask for all neurons (b8). **(C)** Cells identified in the entire field of view using our workflow. **(D)** Strategy for combining cellular fragments into cells based on spatial proximity (Euclidean distance) and activity similarity (temporal cross-correlation). A threshold (<0.1) is applied to the geometric mean of the matrices to determine which fragments to combine.

**Fig. S2.**
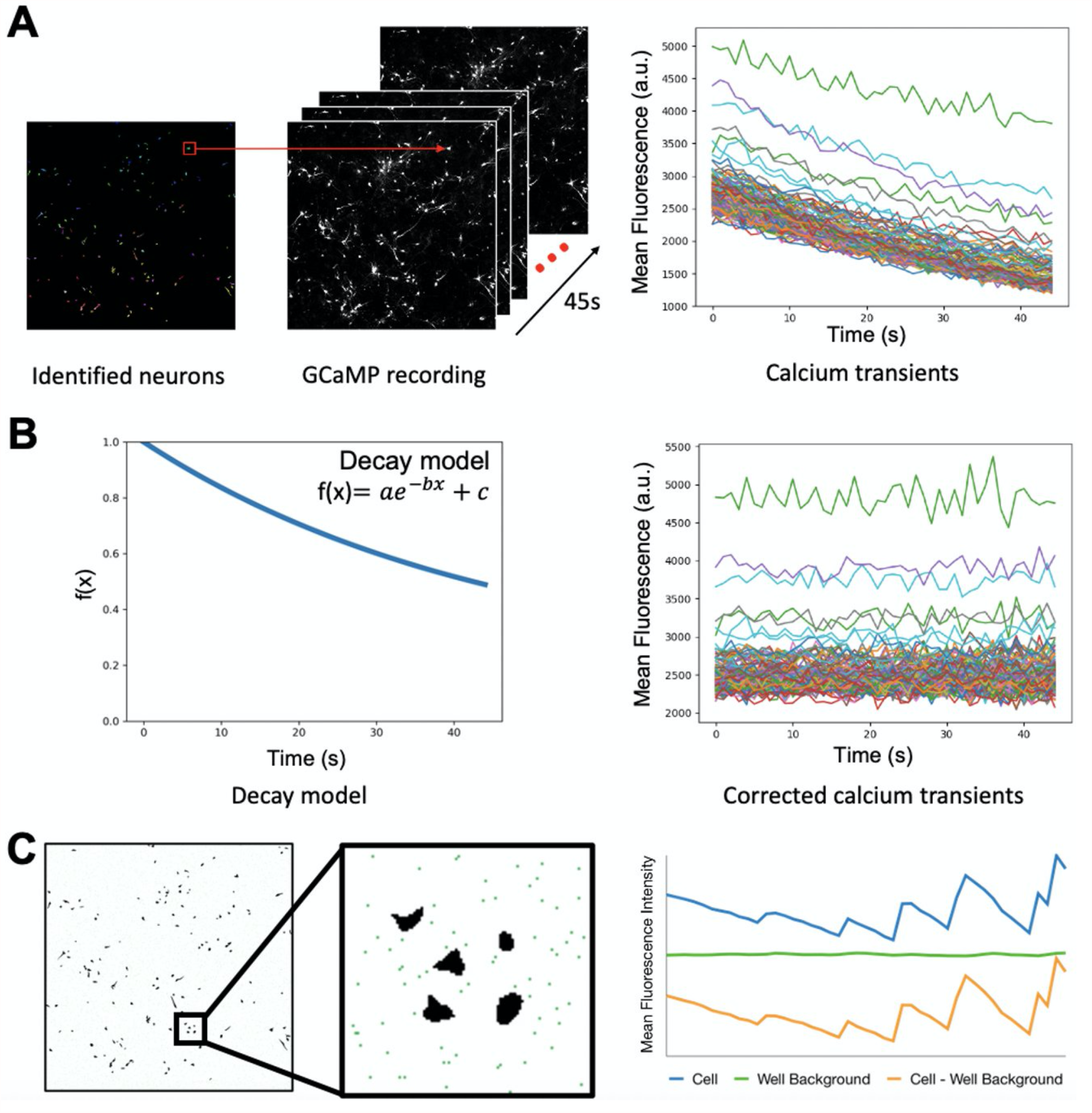
Imaging Artifacts Removal. **(A)** A cell mask is applied to the original GCaMP recording to extract the calcium trace of each spatial footprint. **(B)** Photobleaching effects are modeled as an exponential decay in the GCaMP signal over time and removed from the calcium traces. **(C)** Background luminescence is estimated by averaging the intensity of randomly selected non-cell pixels (green pixels) and removed from a sample cellular calcium trace.

**Fig. S3.**
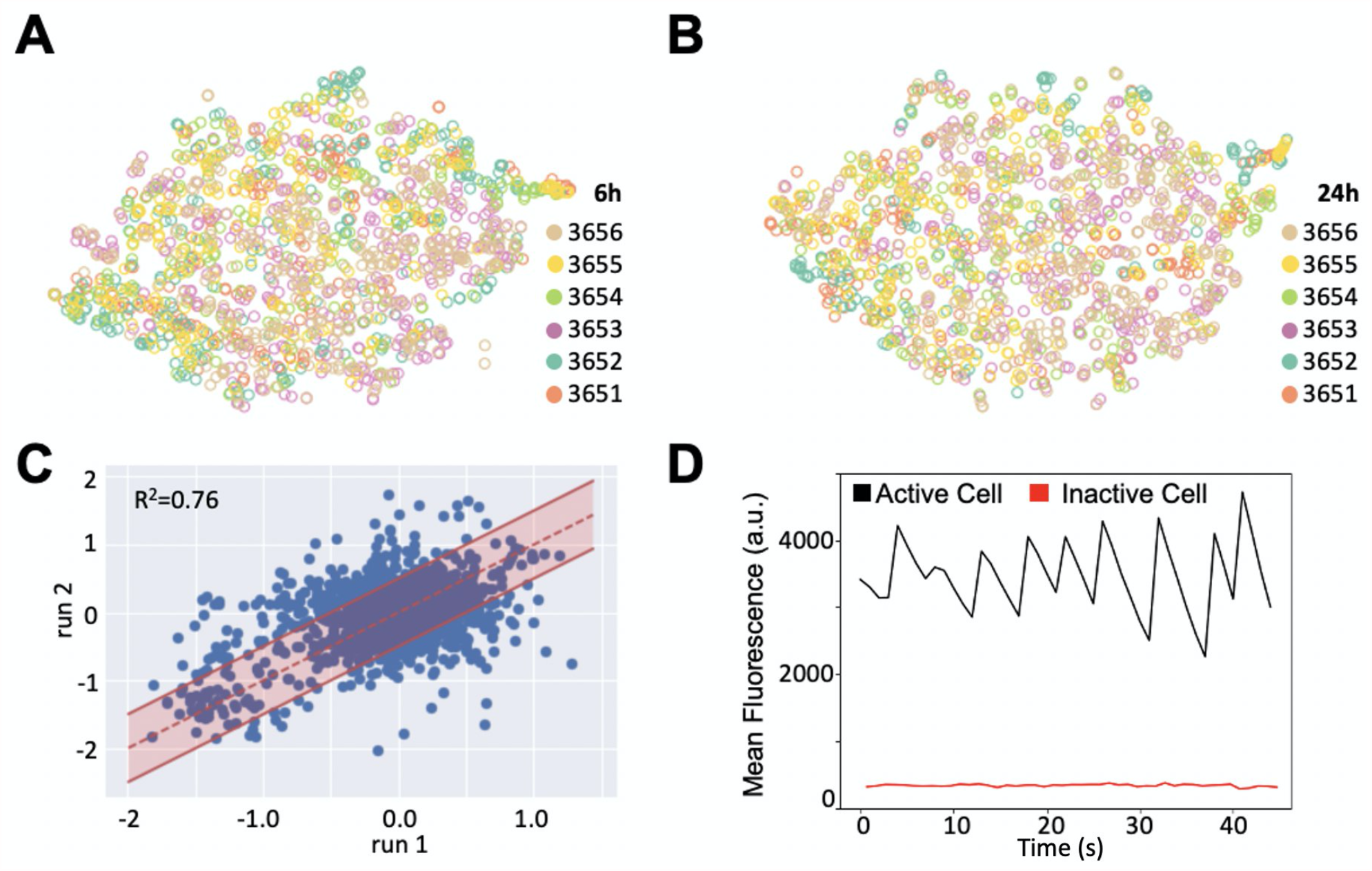
Quality Control. **(A)** Unsupervised dimensionality reduction using t-Distributed Stochastic Neighbor Embedding (t-SNE) shows no plate bias six hours after treatment. **(B)** No plate bias is present even twenty-four hours after treatment. **(C)** Normalized peak counts from two independent recording sessions across all wells show positive correlation between the replicates. A reference correlation line is shown (dashed red line) between ± 1 bands (solid red lines). **(D)** A representative GCaMP signal obtained from an active cell (black) and an inactive cell (red) show separation of the activity states captured by our pipeline.

**Table S1.**
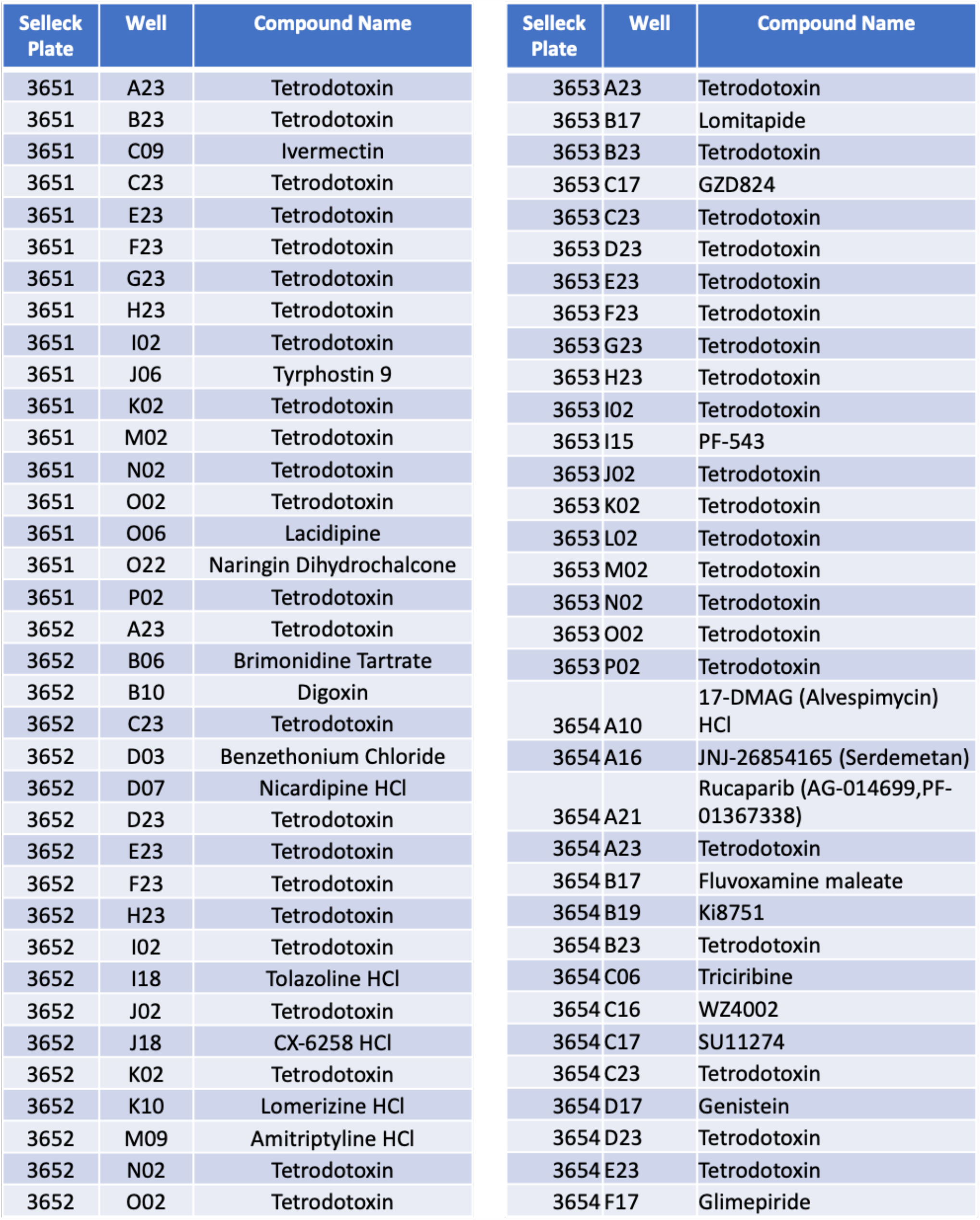
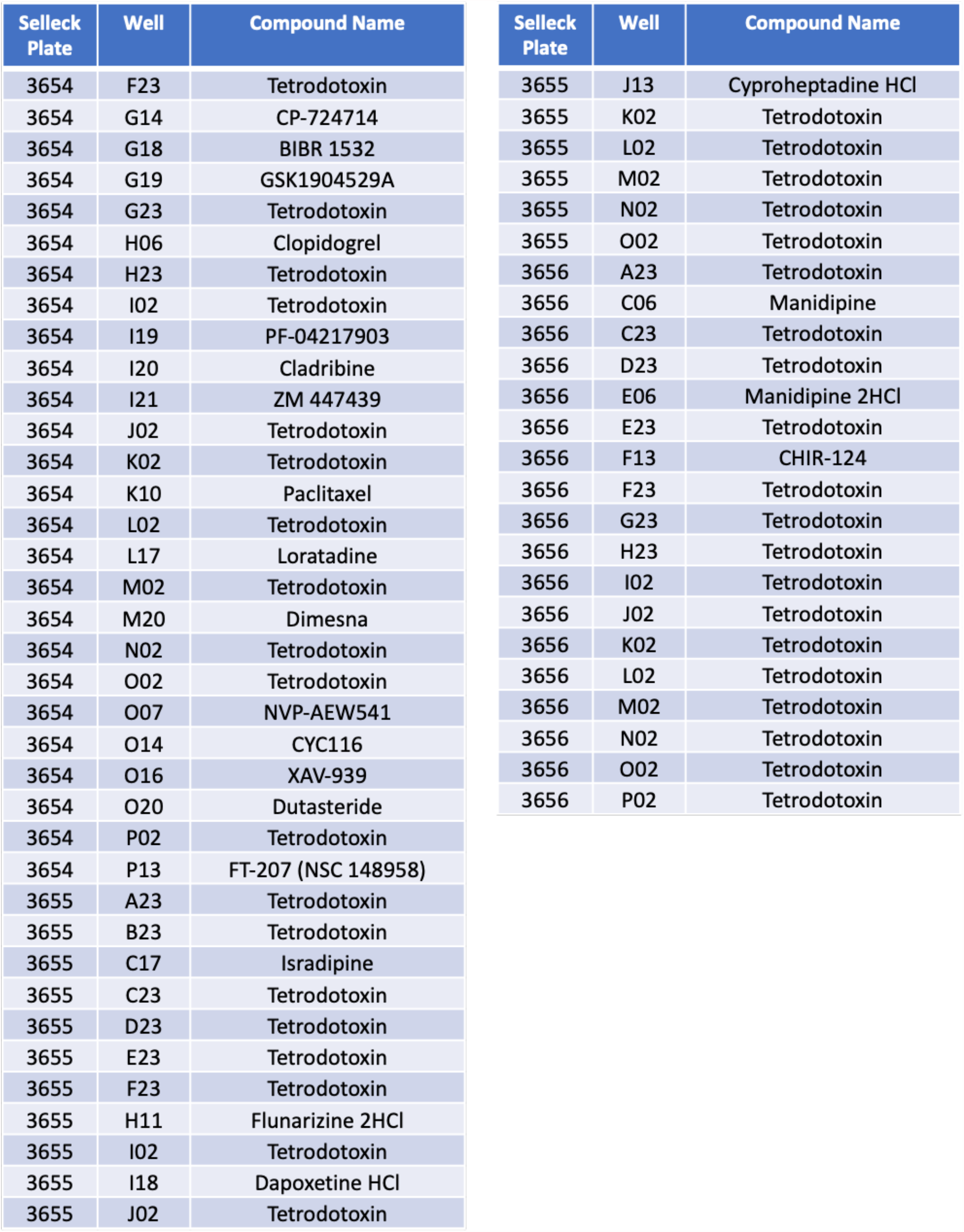
Toxic Compounds. Compounds from our high-throughput screen, including those from the Selleck chemogenomic library, that resulted in low cellular activity (<20 active cells) grouped to show origin plate, well identifier, and compound name.

